# Habenula alterations in resting state functional connectivity among autistic individuals

**DOI:** 10.1101/2025.05.14.653992

**Authors:** Chloe L. Hampson, Julio A. Peraza, Lauren M. Guerrero, Katherine L. Bottenhorn, Michael C. Riedel, Fahad Almuquin, Donisha D. Smith, Katherine M. Schmarder, Katharine E. Crooks, Jaden A. Sangoi, Krystyna R. Keller, Rosario Pintos Lobo, Matthew T. Sutherland, Erica D. Musser, Yael Dai, Rumi Agarwal, Fahad Saeed, Angela R. Laird

## Abstract

**Background:** The reward-based theoretical framework of autism suggests that altered reward circuitry contributes to core symptoms. Recent prior research has revealed autism-related structural alterations in the habenula, a small epithalamic structure associated with motivation and emotion; however, potential alterations in functional connectivity (FC) remain unexplored.

**Methods:** Anatomical and resting state functional magnetic resonance imaging (rs-fMRI) data were accessed for 1,479 participants (*N*=661 autism; *age*_*m*_: 16.68±8.23 years) in the Autism Brain Imaging Data Exchange (ABIDE). To investigate habenula alterations, we conducted a whole-brain resting state FC analysis using manually delineated subject-specific seeds, followed by regression analyses to explore age and brain-behavior interactions.

**Results:** Across the entire sample, extensive habenula connectivity was observed within the midbrain dopaminergic reward system. Compared to neurotypical (NT) controls, autistic participants exhibited significantly increased habenular connectivity with the bilateral middle and superior temporal gyri. From childhood to early adulthood, autistic adolescents displayed an accelerated developmental habenula FC trajectory than NTs with the cingulate gyrus. Between groups, habenula hyperconnectivity was inversely associated with behavioral scores for social motivation and communication.

**Conclusions:** This study provides novel evidence of habenula connectivity alterations in autism, highlighting atypical FC with sensory processing regions. Further findings suggest that habenula circuitry develops differently among autistic adolescents, with links between habenula hyperconnectivity and social behaviors. Taken together, these results contribute to emerging evidence that the dopaminergic reward system may play a critical role in the pathophysiology of autism.

## Introduction

Autism is a lifelong neurodevelopmental disability characterized by social communication differences, as well as restrictive interests and/or repetitive behaviors (RRB) (1). Extending beyond the core criteria, autistic individuals^1^ face everyday challenges in achieving positive life outcomes (5), with previous research highlighting below age-expected levels of executive functioning (EF) (6) and daily living skills (DLS) (7) among autistic adolescents. The social motivation (SM) theory argues that these differences, and challenges, are due to diminished social orienting, social reward, and social maintenance (8), thereby providing a theoretical framework of understanding autism centered on social functioning. This emphasis aligns with many neuroimaging studies that have aimed to elucidate underlying neurobiological mechanisms of autism, yielding a predominance of studies that have largely focused on structural and functional alterations in areas of the “*social brain*” (9). However, to better capture the autistic behavioral spectrum, the social motivation framework was expanded to a reward-based framework that is centered more broadly around constructs related to motivation and affect, thus linking the social differences and RRB that are observed in autism (10,11).

The reward-based theoretical framework suggests that alterations in the brain’s reward circuitry contribute to the core behaviors associated with autism. The reward network, which includes the anterior cingulate cortex, orbital prefrontal cortex, ventral striatum, ventral pallidum, and midbrain dopamine neurons (12), has been found to play a critical role in adaptive motivated behavior (13), as well as guiding both social and nonsocial learning and behavior throughout development (14). Of the regions comprising the reward network, the striatum is repeatedly implicated in autism given its functions that relate to the regulation of behavioral flexibility, motivational state, goal-directed learning, and attention (15). As a key dopaminergic region, the cortico-striatal system shows robust evidence of autism-related alterations in function (16) and structure (17) across development, supporting the reward-based theoretical framework of autism. Dopaminergic pathways within the reward network, including the striatum, are modulated by the habenula, a paired midline structure located on the dorsal surface of the thalamus. Importantly, the functional role of the habenula in autism has yet to be studied.

The habenula is a small epithalamic structure that is divided into two functionally and cytologically distinct medial and lateral nuclei that serve as a connecting link among basal forebrain, striatal, and midbrain regions (18). Task-based animal models have suggested that the habenula plays a substantial role in reward prediction, motivation (19), and aversion processing (20), with replicated involvement in reward prediction in human models (21). To better understand the functional role of the human habenula, neuroimaging studies have mapped resting state habenula connectivity using 3T (22) and 7T functional magnetic resonance imaging (fMRI) (23) among neurotypical (NT) individuals. Torrisi et al. (23) highlighted habenula coupling with forebrain regions that were strongly implicated in emotion and motivation processing, postulating that the habenula plays a role in establishing and maintaining emotional states in humans. Due to its linkage to numerous functional domains relevant to psychopathology, habenula structural and functional connectivity (FC) alterations have been investigated in autism co-occurring behaviors such as those characterized by ADHD (24), major depressive disorder (25), and schizophrenia (26). Although structural alterations in the habenula have been observed in autism (27), potential functional alterations, which may contribute to autistic behaviors relating to emotion and motivation processing, currently remain unexplored.

Addressing this gap in the autism literature, we accessed resting state fMRI data from the Autism Brain Imaging Data Exchange (ABIDE) (28,29) to explore the habenula FC alterations underlying autism. Primarily, we sought to map habenula FC in autistic individuals by conducting a seed-based, whole-brain resting-state FC (rsFC) analysis. Given that differences in striatal volume (30,31) are associated with FC alterations in autism (32), we hypothesized that the known structural differences in the habenula might similarly correspond to atypical habenula FC among autistic individuals compared to NT controls. To provide supplementary insight into autism-related habenula connectivity patterns, we performed two separate regression analyses. In the first regression analysis, we probed for age-related changes in habenula connectivity between autistic individuals and NT controls. Due to recent work revealing an increase in striatal connectivity with cerebellar regions from childhood to early adulthood in autism (33), we hypothesized to observe similar age effects in habenula connectivity. In the second exploratory regression analysis, we aimed to examine the relationship between altered habenula connectivity and autistic behaviors by accessing the phenotypic data available in the ABIDE dataset. To capture the broader domains that influence everyday functioning, we selected phenotypic measures that assessed the level of impairment an individual experienced in SM, social communication (SC), EF, and DLS. As the habenula is primarily implicated in modulating motivation and adaptive behavior, we hypothesized that habenula connectivity would be significantly associated with SM and DLS, but not SC or EF.

## Methods

### Participants

Participants for the current study were selected from the large-scale, multisite ABIDE I and II datasets. The combined ABIDE dataset consisted of neuroimaging data, as well as, demographic and phenotypic information from approximately 2,000 participants across 24 international sites. Written informed consent was obtained through institutional IRB. Further information regarding the ABIDE data collection can be found at http://fcon_1000.projects.nitrc.org/indi/abide/abide_I.html. Demographic information was provided, including age at scan and sex, and autism spectrum disorder^2^ (ASD) diagnoses were made according to the Diagnosis and Statistical Manual of Mental Disorders—4th edition (DSM-IV) (34) and 5th edition (DSM-5) (35).

### Phenotypic Measures

The phenotypic measures included in this study assessed behavioral differences commonly occurring in autism, encompassing SM, SC, EF, and DLS. SM refers to the extent to which an individual is generally motivated to engage in social-interpersonal behavior, which was measured with the 11-item Social Responsiveness Scale (SRS) Social Motivation subscale (36). SC refers to the reciprocal and expressive communication in social situations, which was measured with the 22-item SRS Social Communication subscale (36). EF refers to working memory, flexible thinking, and self-control, which was measured through the Behavior Rating Inventory of Executive Function (BRIEF) Global Executive Composite Score (GEC) T-score (37). The BRIEF GEC T-score measures EF by compiling scores from three composite indexes that test for behavior regulation, emotional recognition, and cognitive regulation. DLS are a measure of adaptive behavior encompassing everyday tasks necessary for independent living, including personal care, household responsibilities, and community activities such as maintaining hygiene, preparing meals, and managing money and time. DLS were measured with the DLS domain of Vineland-2^nd^ edition (38).

### Neuroimaging Data Acquisition and Preprocessing

Resting state, functional, and structural MRI data were acquired across the 24 scanning sites using 3T MRI scanners. For all participants, a T1-weighted structural image was collected and used for registration to Montreal Neurological Institute (MNI) space. Full details for acquisition parameters, informed consent, and site-specific protocols can be found at http://fcon_1000.projects.nitrc.org/indi/abide/abide_I.html. Data were preprocessed using fMRIPrep 23.1.3, which utilizes the Brain Imaging Data Structure (BIDS) to ensure high-quality preprocessing with minimal user intervention (39,40). The anatomical images were intensity-corrected, skull-stripped, segmented, and spatially normalized to a standard brain template in MNI152 space. Furthermore, the preprocessed data underwent both manual and automated quality control. Using the ratings reported by independent raters for ABIDE I, we manually excluded participants that received a “*fail*” rating from any of the raters. Additionally, manual verification was performed for all preprocessed T1-weighted images. We then used MRIQC 23.1.0 to inspect, flag, and remove participants, sessions, or runs (39) if they had excessive head motion defined by a framewise displacement greater than 0.35mm or had less than 100 usable time points following motion censoring (41). Lastly, denoising was performed on the remaining participants using AFNI’s `3dTproject`, which utilizes a linear regression model to nuisance time series from each voxel within a dataset.

### Habenular Region of Interest

Bilateral habenula ROIs were manually delineated on each participant’s preprocessed T1-weighted image in MNI152 space (**Fig. 1**). Using Mango Image Viewer (http://mangoviewer.com), we followed the procedure for manual habenula delineation outlined by Lawson et al. (42). This process yielded unique, subject-specific bilateral habenula ROIs for each participant, reflecting precise individual anatomical variability.

**Figure 1.**
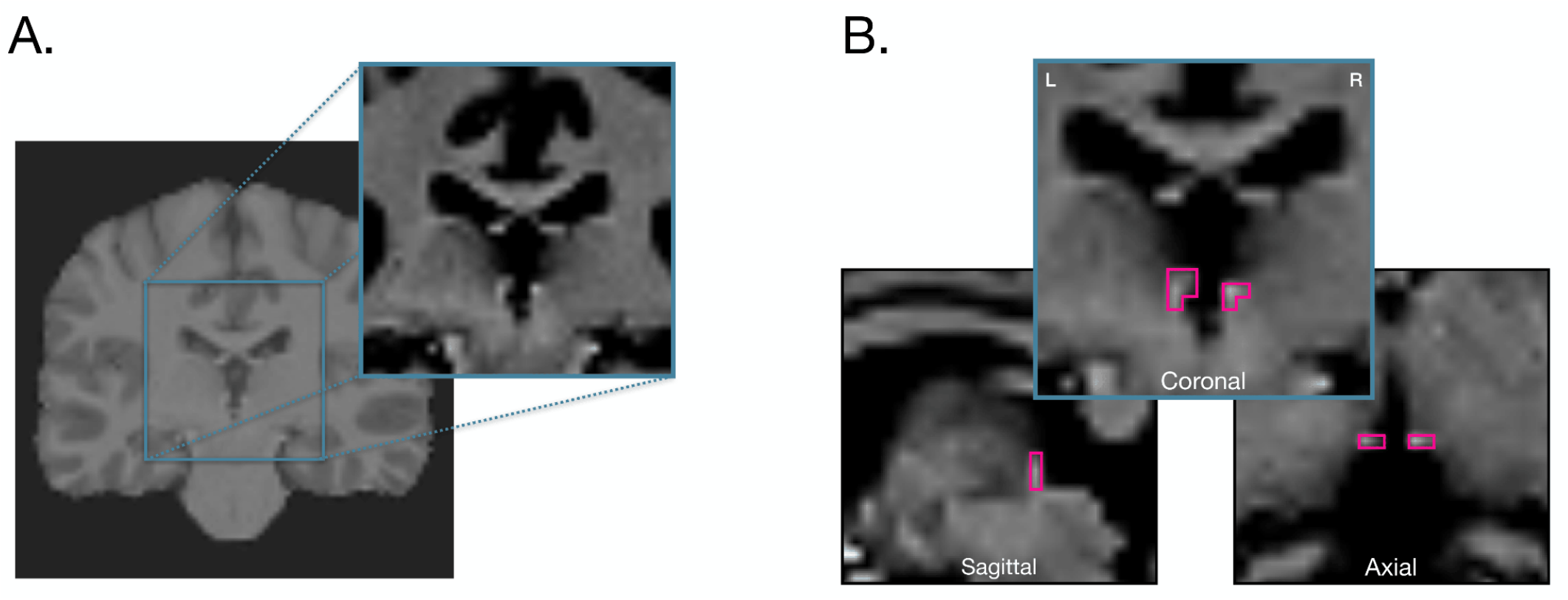
Bilateral Habenula Delineation. Each participant’s preprocessed T1-weighted image was loaded into Mango Image Viewer. **A)** In the coronal view, the midbrain was examined to locate the third ventricle, upon which the image was zoomed in and contrast was increased to improve the visibility of the white matter structures. **B)** Coronal, sagittal and axial views of a manually delineated bilateral habenula region of interest (volume = 64mm^3^).

### Whole-Brain Habenula Functional Connectivity Analysis

#### Group Averaged Habenula Connectivity

Participant-level FC analyses were conducted using AFNI (v22.3.05) (43). Voxelwise time series were extracted from each ROI (i.e., left and right habenula) using the unsmoothed, preprocessed, and denoised data via AFNI’s `3dmaskave`. For each participant, averaged ROI time series were generated by calculating the mean voxel value for each time point across non-zero voxels respective to each region. The averaged ROI time series were entered into a general linear model to generate whole-brain correlation maps for each participant. The resulting z-transformed (Fisher *r*-to-*z* transformation) correlation maps were entered into group-level linear mixed effects models with diagnosis (ASD, NT) as a categorical factor and age as a continuous factor. Significant site-related interaction effects were observed for all covariates, including group, age, and sex (**Figs. S1, S2, S3**); thus, we determined that sex and site would be best controlled with AFNI’s 3dLMEr (44,45), as ComBat may distort site variance when covariates are imbalanced (46). A group-averaged map (ASD+NT) was generated to visualize regions exhibiting positive and negative habenula connectivity for the combined ASD and NT groups. Thresholded z-maps were generated using `3dClustSim` to compute a cluster size at a voxel threshold of *p* = 0.0001, and a cluster threshold of *p* = 0.01.

#### Group Differences in Habenula Connectivity

The group-averaged map was used to mask regions functionally connected with the habenula in both the ASD and NT groups. A group-difference map was generated to identify regions that exhibited atypical habenula rsFC among ASD participants (ASD vs. NT). Thresholded z-maps were generated at a voxel threshold of *p* = 0.0001 and a cluster threshold of *p* = 0.01.

### Age-Related Variability in Habenula Connectivity

Next, a whole-brain regression analysis was performed to probe for age-related changes in habenula connectivity. Previous studies have identified substantial rsFC variability from childhood to early adulthood, characterized by a refinement of functional networks in the brain (47). Thus, we chose to focus on participants aged 5-21 years, which included the majority of ABIDE participants. For the age-restricted subset of participants, the z-transformed participant-level correlation maps were entered into a second linear mixed effects model in AFNI to examine age-by-group interactions, controlling for sex and scanning site. Thresholded z-maps were generated with a voxel threshold of *p* = 0.005 and a cluster threshold of *p* = 0.05. Clusters exhibiting significant age interactions in habenula connectivity between groups were identified and used to create binarized masks to extract z-scored beta coefficients. For each of the clusters, beta coefficients were averaged and entered into a linear regression model to determine the direction of developmental changes in connectivity for each group, following the approach outlined by Padmanabhan et al. (33).

### Functional Decoding of Habenula Connectivity

We performed functional decoding (48) on each of the thresholded z-maps to provide additional information regarding the functional role of the habenula among ASD participants. Specifically, decoding was conducted to characterize the functional role of the regions identified in each step of our analysis. To perform functional decoding, we utilized `Gradec` (v0.0.1rc5), a Python package designed for meta-analytic functional decoding (49). Within this package, the decoder algorithms utilize the Neurosynth database to conduct large-scale, automated synthesis of fMRI data (50). Currently, the Neurosynth has over 500,000 activation coordinates from 14,371 task-based neuroimaging studies (50). Using `Gradec`, we extracted Neurosynth terms that were correlated with the habenula connectivity maps and generated word clouds and radar plots to visualize the correlations between the terms and the spatial distribution of connectivity (49).

### Associations Between Habenula Connectivity and Autism Symptomatology

Lastly, a separate regression analysis was performed to examine associations between atypical habenula connectivity and phenotypic measures, including SM, SC, EF, and DLS. Matching the approach used in the age-related regression analysis, z-scored beta coefficients were extracted from clusters showing atypical habenula connectivity using a binarized cluster mask. Given that fewer participants had scores for each phenotypic measure, we tested clusters for significant phenotypic main effects among ASD participants, as well as phenotypic interactions between ASD and NT groups (*p* < 0.05) using group as a categorical factor and age as a continuous factor, while controlling for sex and scanning site. For clusters that were identified to exhibit a significant effect, correlation coefficients were entered into a linear regression model as a function of subscore for each of the phenotypic measures. To correct for multiple comparisons, the Benjamini-Hochberg procedure was applied to control the false discovery rate (*p* < 0.05) (51).

## Results

### Sample Selection

From ABIDE participants who reported complete baseline demographic information, we excluded participants from the current study if their MRI data did not meet the previously defined inclusion criteria. The final sample included 1,479 participants (*age*_*m*_: 16.49 ± 8.15 years) from the ABIDE I and II datasets, with 661 ASD participants (**Table 1**). Among the included participants, only a fraction completed the phenotypic measures; therefore, we accessed SM and SC scores for 279 ASD and 330 NT, EF scores for 133 ASD and 200 NT, and DLS scores for 102 ASD and 68 NT. Of the included demographic variables (i.e., age, sex, handedness, diagnostic category, comorbidity, and medication status), significant differences (*p* < 0.001) between the ASD and NT groups were found for sex, diagnostic category, and medication status.

**Table 1.** Participant Information.

|  | ASD (N = 661) |  | NT (N = 879) |  | t-value | p-value |
| --- | --- | --- | --- | --- | --- | --- |
|  | M | SD | M | SD |  |  |
| Age (in years) | 16.68 | 8.23 | 16.34 | 8.08 | 0.815 | 0.415 |
| | N | | N | | $\chi^2$ -value | p-value |
| Sex | -- |  | -- |  | 31.29 | < 0.001 |
| Male | 572 |  | 611 |  | -- | -- |
| Female | 89 |  | 207 |  | -- | -- |
| Handedness | -- |  | -- |  | 9.90 | 0.007 |
| Right | 450 |  | 613 |  | -- | -- |
| Left | 55 |  | 39 |  | -- | -- |
| Mixed | 23 |  | 23 |  | -- | -- |
| Diagnostic Category | -- |  | -- |  | 685.9 | < 0.001 |
| Control | 12 |  | 405 |  | -- | -- |
| Autism | 222 |  | 0 |  | -- | -- |
| Asperger's | 66 |  | 0 |  | -- | -- |
| PDD-NOS | 28 |  | 0 |  | -- | -- |
| Comorbidity | -- |  | -- |  | -- | -- |
| ADHD | 7 |  | 0 |  | -- | -- |
| Anxiety | 18 |  | 0 |  | -- | -- |
| Disruptive Disorder | 2 |  | 0 |  | -- | -- |
| Mood Disorder | 7 |  | 0 |  | -- | -- |
| Medication Status | -- |  | -- |  | 192.6 | < 0.001 |
| Medicated | 164 |  | 12 |  | -- | -- |
| Not Medicated | 379 |  | 657 |  | -- | -- |

Note: Values are provided as mean (M) ± standard deviation (SD). t-value was obtained using a two-sample t-test. $\chi^2$ values were obtained using chi-squared tests. < 0.001 indicates significant differences between the ASD and NT groups for demographic variables. Abbreviations: ASD, autism spectrum disorder; ADHD, attention deficit hyperactivity disorder; M, mean; NT, neurotypical; PPD-NOS, pervasive developmental disorder-not otherwise specified; SD, standard deviation.

### Whole-Brain Habenula Functional Connectivity Analysis

#### Group-Averaged Habenula Connectivity

The whole-brain, group-averaged (ASD+NT) habenula connectivity map (**Fig. 2A**) depicts positive and negative habenula connectivity through an extensive network of bilateral cortical and subcortical regions (**Table 2**). Neurosynth-based functional decoding revealed that this connectivity pattern is commonly associated with terms for fear, reward, and emotion processing (**Fig. 2C**).

**Table 2.** Peak coordinates for clusters of positive and negative group-averaged (ASD+NT) habenula connectivity, controlling for age, sex, and scanning site.

| X | Y | Z | Peak Stat | Cluster Size | Region |
| --- | --- | --- | --- | --- | --- |
| **Positive Habenula Connectivity** |  |  |  |  |  |
| 4 | -24 | 4 | 27.47 | 542299 | Thalamus |
| -24 | 2 | 68 | 6.07 | 2366 | Left Superior Frontal Gyrus (BA 6) |
| 40 | 4 | 48 | 5.75 | 2546 | Right Middle Frontal Gyrus (BA 6) |
| -31 | -56 | 46 | 5.54 | 1570 | Left Superior Parietal Lobe (BA 7) |
| 48 | -12 | 48 | 5.17 | 690 | Right Precentral Gyrus (BA 4) |
| -46 | -44 | 42 | 5.15 | 648 | Left Inferior Parietal Lobe |
| 60 | 10 | 12 | 5.09 | 901 | Right Inferior Frontal Gyrus (BA 44) |
| -40 | -20 | 42 | 5.04 | 522 | Left Postcentral Gyrus (BA 2) |
| -40 | -78 | 33 | 5.01 | 1048 | Left Superior Occipital Gyrus (BA 19) |
| -22 | -30 | 68 | 4.96 | 645 | Left Postcentral Gyrus (BA 3) |
| -28 | 54 | 16 | 4.95 | 2345 | Left Superior Frontal Gyrus (BA 10) |
| 28 | 28 | 41 | 4.76 | 1355 | Right Middle Frontal Gyrus (BA 8) |
| -38 | -4 | 56 | 4.73 | 654 | Precentral Gyrus (BA 6) |
| 20 | -29 | 73 | 4.55 | 610 | Right Postcentral Gyrus (BA 3) |
| -30 | 30 | 39 | 4.52 | 3744 | Left Middle Frontal Gyrus (BA 8) |
| **Negative Habenula Connectivity** |  |  |  |  |  |
| -20 | -10 | 29 | -6.31 | 3096 | Left Caudate |
| 32 | -42 | 19 | -5.67 | 1981 | Right Insula (BA 13) |
| 18 | 32 | 7 | -5.60 | 4686 | Right Anterior Cingulate (BA 32) |
| -20 | 32 | 9 | -5.14 | 1319 | Left Anterior Cingulate (BA 32) |

**Figure 2.**
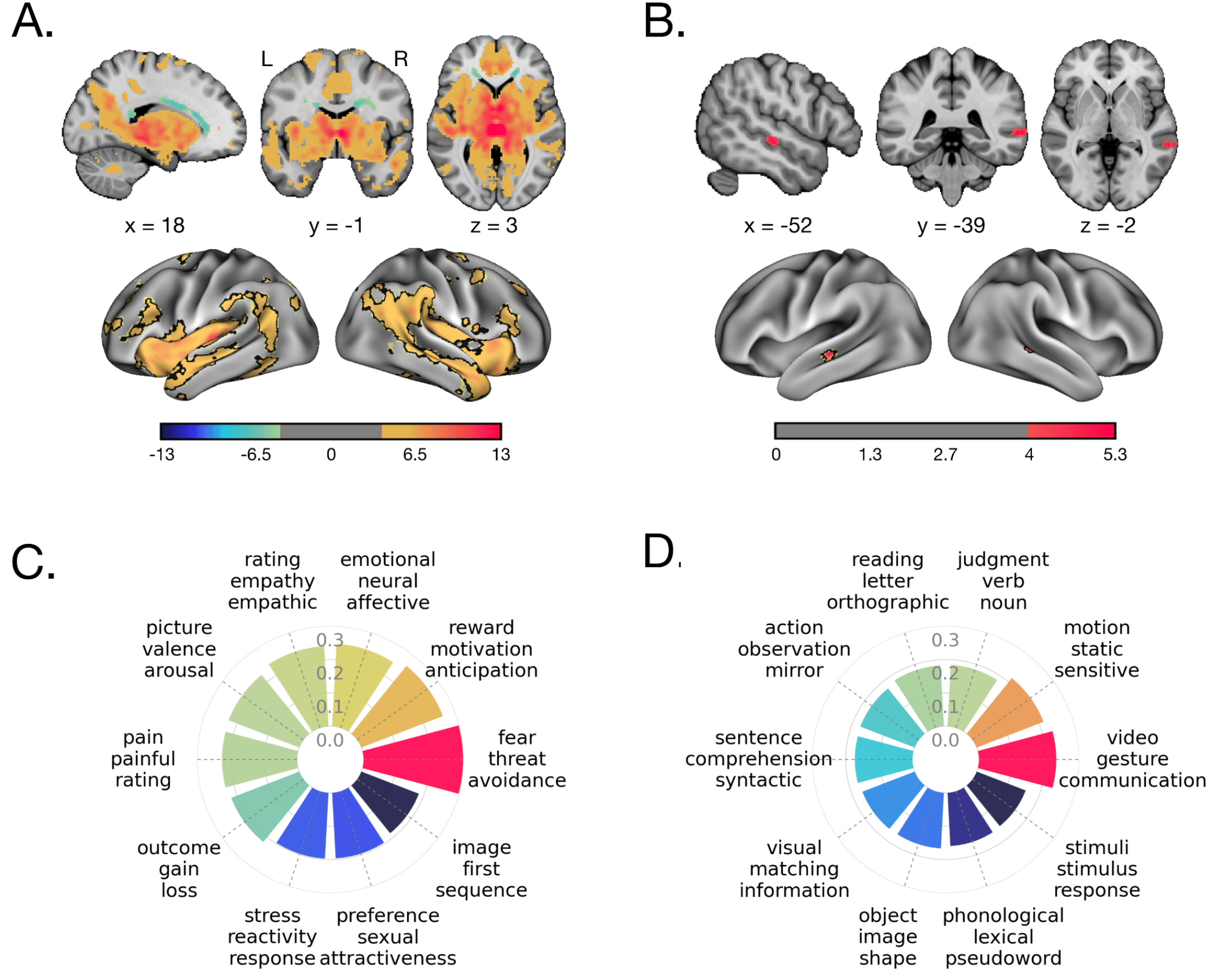
Whole-Brain Habenula Connectivity. Thresholded z-maps and Cohen’s *d* effect maps were generated using 3dClustSim to compute a cluster size at a voxel threshold of *p* = 0.0001 and a cluster threshold of *p* = 0.01. **A)** For the group-averaged (ASD+NT) habenula connectivity, warm colors denote areas of positive connectivity with the habenula and cool colors denote areas of negative connectivity. **B)** For the group-difference (ASD>NT) habenula connectivity, warm colors denote areas of increased connectivity with the habenula among ASD participants. Functional decoding results for the habenula connectivity z-maps are shown as a radar plot with **C)** for group-averaged (ASD+NT) habenula connectivity and **D)** group-difference (ASD>NT) habenula connectivity.

#### Group Differences in Habenula Connectivity

The whole-brain group-difference (ASD>NT) habenula connectivity map (**Fig. 2B**) depicts two regions of *increased* habenula connectivity for the ASD group (**Table 3**). Connectivity was observed with peaks in the bilateral middle temporal gyri (MTG) that extended through the bilateral superior temporal gyrus (STG). Neurosynth-based functional decoding revealed that these regions of increased habenula connectivity among ASD participants are commonly associated with terms for visual, auditory, and action processing (**Fig. 2D**). No statistically significant regions of *decreased* ASD connectivity were observed.

**Table 3.** Peak coordinates for clusters of increased habenula connectivity among ASD participants compared to neurotypical controls (ASD>NT), controlling for age, sex, and scanning site.

| X | Y | Z | Peak Stat | Cluster Size | Region |
| --- | --- | --- | --- | --- | --- |
| 64 | -38 | 1 | 5.22 | 885 | Right Middle Temporal Gyrus (BA 21) |
| -52 | -22 | -2 | 4.92 | 902 | Left Middle Temporal Gyrus (BA 21) |

### Age-Related Variability in Habenula Connectivity

Next, we probed for age-related variability from childhood to early adulthood in habenula connectivity among participants between the ages of 5 and 21 years. The subset sample included 1,145 participants (*N*=516 ASD, 639 NT). The age-related variability analysis identified altered developmental trajectories in habenula connectivity between the ASD and NT groups with the cingulate gyrus (BA 31, **Table 4**). Neurosynth-based functional decoding revealed that the region exhibiting age-related variability in habenula connectivity is commonly associated with terms related to decision making and memory (**Fig. 3A**). Entering the averaged z-scored beta coefficients into a linear regression model revealed increasing yet deviating developmental trajectories of habenula connectivity with the cingulate gyrus, with ASD having a steeper slope than the NTs (Cohen’s *d* = 0.076, **Fig. 3B**). Although AFNI identified this region as showing a significant age-by-group interaction, averaging beta coefficients across the cluster likely diluted spatially localized effects, weakening the interaction in the regression analysis.

**Table 4.** Peak coordinates for clusters of regions that showed an age by group interaction, controlling for sex and scanning site.

| X | Y | Z | Peak Stat | Cluster Size | Region |
| --- | --- | --- | --- | --- | --- |
| 4 | -34 | 40 | 5.36 | 3,432 | Cingulate Gyrus (BA 31) |

**Figure 3.**
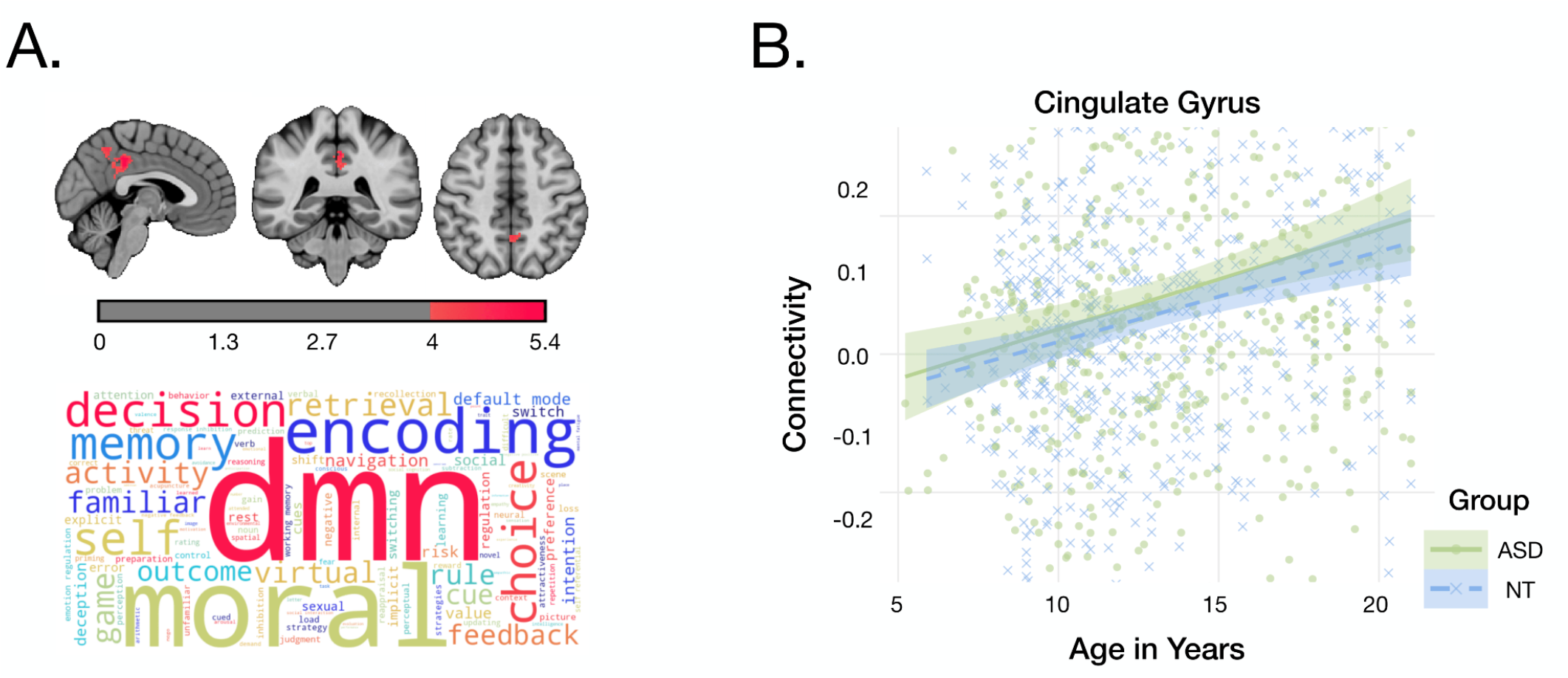
Age-Related Variability in Habenula Connectivity. Thresholded z-maps and Cohen’s *d* effect maps were generated with a voxel threshold of *p* = 0.005 and a cluster threshold of *p* = 0.05. Warm colors denote areas of increased connectivity with the habenula. **A)** Volumetric brain slices (top) depict regions with age-related variability in habenula connectivity with functional decoding results shown as a word cloud (bottom). **B)** Z-scored beta coefficients from the cingulate gyrus were entered into a linear regression model as a function of age to determine differences in developmental slope between the ASD (solid green) and NT (dashed blue) groups. For effect sizes between groups, the rMTG showed a small positive effect (*d* = 0.076).

### Associations Between Habenula Connectivity and Autism Symptomatology

Given the small sample sizes with completed phenotypic information, we note the following analyses were considered exploratory. For the two regions that exhibited altered habenula connectivity between the ASD and NT groups (**Table 3**); z-scored beta coefficients were extracted and entered into a linear regression model to test for significant (*p* < 0.05) main and between-group interaction effects (**Table 5**) for each of the phenotypic measures. For SM and SC, a significant interaction was identified in the rMTG. For both social phenotypes, higher behavior scores were associated with slightly increasing habenula connectivity in the ASD group, whereas the NT group showed increasing connectivity (**Fig. 4**). No significant main effects for the ASD group were observed. For EF and DLS, no significant main or interaction effects were found. Despite these findings, none survived the correction for multiple comparisons.

**Table 5.** Group-difference (ASD>NT) main effect and interaction of phenotypic measures on habenula connectivity.

| Region | Group Interaction |  |  |  | ASD Main Effect |  |  |  |
| --- | --- | --- | --- | --- | --- | --- | --- | --- |
|  | Estimate | SE | p-value | p<sub>FDR</sub>-value | Estimate | SE | p-value | p<sub>FDR</sub>-value |
| **Social Motivation** |  |  |  |  |  |  |  |  |
| **rMTG** | 0.0075 | 0.00302 | **0.0133** | 0.0687 | <0.001 | 0.00168 | 0.834 | 0.96 |
| **IMTG** | 0.00376 | 0.00251 | 0.135 | 0.270 | -0.00251 | 0.00139 | 0.0714 | 0.571 |
| **Social Communication** |  |  |  |  |  |  |  |  |
| **rMTG** | 0.00449 | 0.00188 | **0.0172** | 0.0687 | <0.001 | <0.001 | 0.960 | 0.960 |
| **IMTG** | <0.001 | 0.00157 | 0.606 | 0.802 | <0.001 | <0.001 | 0.676 | 0.960 |
| **Executive Functioning** |  |  |  |  |  |  |  |  |
| **rMTG** | <0.001 | 0.00145 | 0.77 | 0.802 | <0.001 | <0.001 | 0.931 | 0.960 |
| **IMTG** | -0.00152 | 0.00133 | 0.255 | 0.409 | <0.001 | <0.001 | 0.925 | 0.960 |
| **Daily Living Skills** |  |  |  |  |  |  |  |  |
| **rMTG** | <0.001 | 0.00224 | 0.802 | 0.802 | 0.00157 | 0.00161 | 0.332 | 0.885 |
| **IMTG** | 0.00314 | 0.00193 | 0.106 | 0.270 | -0.00169 | 0.00134 | 0.209 | 0.836 |

Note: Main effects and interactions were significant at $P < 0.05$ when controlling for age, sex, and scanning site.
Abbreviations: BA, Brodmann area; FDR, false discovery rate; MTG, Middle Temporal Gyrus; SE, standard error.

**Figure 4.**
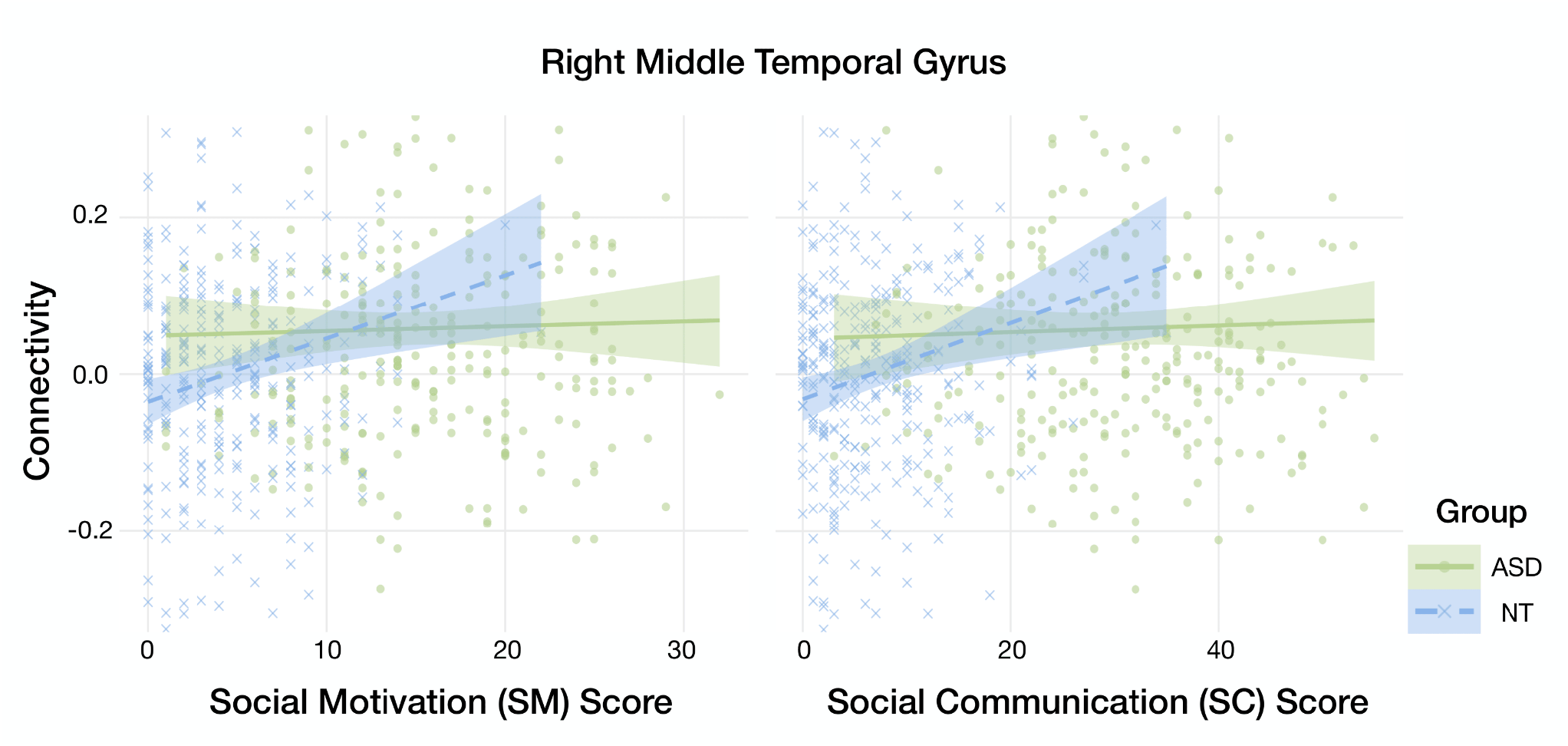
Associations Between Habenula Connectivity and ASD Symptomatology. Z-scored beta coefficients were extracted from regions that displayed increased habenula connectivity for the ASD group and entered into a linear regression model to test for significant phenotypic main and interaction effects (*p* < 0.05). The regression lines for the ASD (solid green) and NT (dashed blue) groups denote the relationship between the phenotypic score and connectivity for each region. For SM and SC scores, an interaction effect was found in the rMTG.

## Discussion

We investigated resting state functional connectivity (rsFC) of the habenula among autistic individuals. Across the entire sample, group-averaged (ASD+NT) results confirmed existing evidence that the human habenula functionally interacts with the midbrain dopaminergic reward system (52). Group-difference (ASD>NT) analyses identified atypical habenula hyperconnectivity in autism within bilateral middle temporal gyri (MTG). Further investigation of variability through age-related regression analyses revealed alterations in connectivity from childhood to early adulthood (ages 5–21) between autistic individuals and NTs, particularly with the cingulate gyrus (BA 31). Finally, a second exploratory regression analysis demonstrated that atypical hyperconnectivity in autism had a significant interaction effect with social motivation (SM) and social communication (SC), but not executive function (EF) and daily living skills (DLS). However, due to the small sample of individuals with completed phenotypic information, these results did not survive multiple comparison corrections.

### Whole-Brain Habenula Functional Connectivity: ASD+NT and ASD>NT

Our study identified widespread habenula connectivity among the entire sample in task-positive brain regions (53). Specifically, we report prominent positive habenula connectivity with key subcortical regions of the reward network, such as the thalamus, striatum, and several midline structures, aligning with previous studies (22,23,54). Moreover, we replicated strong positive cortical habenula connectivity with the dorsolateral prefrontal cortex (dlPFC; BA 9), as well as across primary and associative somatosensory cortices. Positive connectivity was also observed with the amygdala, in agreement with Ely et al. (22) and supported by the amygdala’s role in regulating innate and learned behaviors (55). While other previous studies conducted at 3T (54) and 7T (23) reported no significant connectivity between the human habenula and amygdala, we note that the work of Ely et al. (22) and the present study included a more robust sample size. Finally, negative connectivity was observed in a few regions of the default mode network, most prominently with the anterior cingulate cortex. Overall, the patterns of resting state connectivity identified in the current study are consistent with the most recent habenular literature. Extending this, functional decoding highlighted the habenula’s involvement in reward and aversion processes and indicated associations with multiple stimulus types, suggesting that the habenula may serve as a hub for the integration of aversive and appetitive sensory signals. In line with this, animal-based studies have shown that the lateral habenula supports a bottom-up multimodal sensory pathway and produces concurrent emotional effects and motor responses that allow animals to efficiently avoid unfavorable environments (56). In humans, the habenula is speculated to play a role in low-level processing of sensory stimuli by receiving early, low-latency information from multiple sources that rapidly drops off as information propagates through higher sensory processing areas (22). While our decoding results align with findings from Torrisi and Ely, they also provide important additional insights into the sensory processing functions of the habenula.

Among autistic individuals, there is an understanding that atypical sensory perception is a fundamental characteristic and core diagnostic symptom of the disorder (35). Autism-related neuroimaging research has extended this with evidence indicating that atypical sensory traits are rooted in low-level processing across sensory modalities and during multimodal perception in primary sensory cortices (57). Our results revealed atypical habenula hyperconnectivity in autism that peaked in the bilateral MTG and extended through the STG. Consistent with the observed functional differences, prior studies have reported increased gray matter volume in the MTG and STG in autism (58). Functionally, these lateral temporal regions, which include Wernicke’s area, collectively support auditory and language processing, with distinct contributions to speech perception, phonological processing, and semantic integration (59–61). Despite these areas being largely linked to auditory related processes, previous studies have also shown the bilateral STG regions participate low-level, multi-sensory processing, such as the integration of auditory and visual signals (62,63), as well as temporal discrimination of tactile stimuli (64) and processing of olfactory information (65). Our functional decoding results provide further evidence that habenula hyperconnectivity in autism is implicated in multimodal sensory processing, with an emphasis on auditory and visual processes. Given these findings, we posit that habenula connectivity alterations with low-level sensory processing regions may contribute to atypical sensory processing experienced among autistic individuals.

### Age-Related Variability in Habenula Connectivity

Throughout childhood and adolescence, the brain undergoes a multifaceted and dynamic maturation process, wherein alterations of the brain’s development are hypothesized to contribute to autistic behaviors. Although the reorganization of reward circuitry during adolescence is integral to human development (66), developmental alterations of the reward network in autism have only been studied in the striatum. Our results contribute to characterizing age-related variability of the reward network in autism by revealing significant age-related effects on habenula connectivity peaking in the cingulate gyrus, including the posterior cingulate cortex (PCC), and extending through the superior parietal lobule (SPL). Functional decoding of habenular age-related effects observed revealed terms related to decision-making and memories, suggesting that these alterations may reflect differences in memory-guided decision making during reward-seeking. Consistent with this, behavioral studies have demonstrated that while autistic youth may not perform differently in decision-making tasks, their choices are often driven by a motivation to avoid potential losses rather than to seek possible rewards (67). Therefore, considering the lateral habenula’s inhibitory role in predicting imminent aversive events (68), it is possible age-related variability in habenula connectivity in autism could underlie these behaviors.

### Associations Between Habenula Connectivity and Autism Symptomatology

A central challenge in autism research has been to link the pathophysiology of autism with its complex and heterogeneous behaviors. Here, we provide evidence that the habenula links to core domains of social functioning in autism. We observed a significant association with SM in autistic individuals compared to NTs, as hypothesized, as well as an association with SC, although this was not hypothesized. In agreement with previous evidence of the habenula guiding animal social behaviors (69,70), our findings suggest the human habenula plays a role in the motivation to engage in social interactions, as well as the reciprocal and expressive aspects of communication. In indicating that SM and SC are related constructs with similar relationships to reward-related processing in autism, our results are consistent with the SM theory (8), which states that all social behaviors are derived from SM. Although the habenula plays a well-known role in adaptive behaviors, such as modulating responses to changing environments and to aversive and rewarding stimuli (71), we did not include a direct measure of adaptive behavior in the current study. Instead, we examined EF and DLS as proxy measures, neither of which showed an association with habenula connectivity. Therefore, although our group-averaged FC results highlight the central role of the habenula in motivated and adaptive behaviors, which are crucial for positive life outcomes and pose challenges for autistic individuals in achieving independence (72), these findings suggest that altered habenula connectivity underlies behavioral differences in social behavior in autism.

### Limitations and Future Directions

Several key limitations of the present study should be noted. First, due to the small size of the habenula and our standard-resolution functional data, we did not attempt to parcellate the habenula ROI into medial and lateral portions. Although the lateral and medial habenular systems are largely separate from each other (73), a smaller ROI would likely have led to increased signal contamination from surrounding regions. More specifically, as the left and right ROIs contained varying ratios of “*true*” habenula to non-neuronal tissue, which leads to discrepancies in the BOLD signal contamination. Thus, results would be more susceptible to partial voluming effects, reducing any meaningful conclusion that could have been made about their distinct functions. Second, our analysis of age-related variability relied on cross-sectional data from individuals at different developmental stages. While these findings provide valuable insights into connectivity across adolescence, a longitudinal design would offer a more robust understanding of how these neural trajectories evolve over time in autism. Third, as the ABIDE dataset is a multisite consortia, there was variability in scanning protocols, which contributed to discrepancies in collected demographic information and limited availability of phenotypic data. Particularly of interest, data collection of the groups were imbalanced with some sites scanning more NT than autistic individuals. While we accounted for site-related variance in our analysis, future studies should aim for more controlled data collection to minimize these confounds. Lastly, although our results linking habenula connectivity with autism symptomatology are noteworthy, they did not survive multiple comparison correction. While this does not invalidate our findings, it does highlight the need for replication in datasets with larger phenotypic data sample sizes are needed to ensure the robustness of these associations. Despite these limitations, our results remain interesting and highlight the potential importance in targeting the dopaminergic pathways in autism as they may provide a more unified approach to addressing disruptions in motivated and adaptive functioning. Future studies should incorporate repeated measures within individuals to track habenula connectivity changes across development and further investigate FC alterations of other dopaminergic reward regions.

## Acknowledgments

Primary funding for this project was provided by the FIU Embrace Center for Advancing Inclusive Communities, the FIU Office of Research & Economic Development, and the FIU Undergraduate to Graduate Program. Contributions from co-authors were provided with support from NIH R01-MH09606 (JAP, ARL), U01-DA041156 (ARL), U54-MD012393 (RA), and NIH R35GM153434. This project was also supported by funding provided to each of the contributing ABIDE research institutions, which are listed for each site at https://fcon_1000.projects.nitrc.org/indi/abide/.

We thank the undergraduate research assistants in the FIU Neuroinformatics and Brain Connectivity Lab who contributed to the manual delineation and quality control of habenula regions of interest, including Li-Ann Allen, Liberty Allen, Elizbel Carvajal, Amaya Cosio Gonzalez, Brianna Diaz, Natalia Daher, Ivette Cantillo Hernandez, Kayanna Green, Eulogia Omane-Achamfour, Natalie Ryan, Maria Alejandra, and Lane Stevens. Additional thanks to the FIU Instructional and Research Computing Center (IRCC, http://ircc.fiu.edu) for providing the HPC and computing resources that contributed to the research results reported within this paper.

## Data and Code Availability

The neuroimaging and phenotypic data included in this project can be found in the publicly available ABIDE dataset (https://fcon_1000.projects.nitrc.org/indi/abide/). The code for the analyses conducted in this project can be found on Github (https://github.com/NBCLab/abide-analysis).

## Author Contributions

ARL, CLH conceived and designed the project. CLH, JAP, FA and FS preprocessed data. CLH, JAS, KRK contributed to the manual delineation and quality control of habenula regions of interest. CLH, JAP analyzed data, and CLH, ARL, KEC interpreted data. JAP, KLB, MCR, DDS contributed scripts and pipelines. CLH, ARL wrote the paper and all authors contributed to the revisions and approved the final version.

## Footnotes

1 In the current manuscript, we used identity-first language, as recommended by Toboas et. al. (2), Bury et. al. (3), and Kenny et al. (4).

2 When discussing data accessed from ABIDE, we use the term ‘ASD’ when referring to autism to align with the dataset’s terminology.

