## Supplemental Information for "Habenula alterations in resting state functional connectivity among autistic individuals"

### Supplemental Methods

#### Participants

Participants with an autism spectrum disorder (ASD) diagnosis were recruited primarily through the Southwest Autism Resource and Research Center, while other participants with ASD and all age-matched neurotypical (NT) participants were recruited from IRB approved flyers, word of mouth, media, and local support groups. Written informed consent was obtained through institutional IRB. ASD diagnostic procedures included evaluation of an individual's developmental history, clinical observations, and cognitive status by a reliable researcher. To meet the Autism Brain Imaging Data Exchange (ABIDE) inclusion criteria, the diagnosed individual must score above a cutoff in the Autism Diagnosis Observation Schedule—1st (ADOS) (1) or 2nd edition (ADOS-2) (2). NT controls completed the Social Responsiveness Scale—1st (SRS) (3) or 2nd edition (SRS-2) (4) self-report form to screen for significant autism symptoms (anyone with total T score < 60 was included) (5). In addition to the baseline participant demographic information, scanning sites optionally administered additional phenotypic measures to both parents and participants.

#### Phenotypic Measures

Social motivation (SM) was measured with the Social Responsiveness Scale (SRS) Social Motivation subscale, which includes 11 items relating to social disinterest, avoidance of social interactions, and discomfort with interactions (6). For the ASD group, the SRS was administered to a parent for child participants or an informant (parent, a relative, or a significant other), when available, for adult participants. The BRIEF GEC T-score measures executive functioning (EF) by compiling scores from three composite indexes that test for behavior regulation, emotional recognition, and cognitive regulation (7). For the ASD group, the BRIEF was administered to a parent for child participants or an informant (parent, a relative, or a significant other), when available, for adult participants. Daily living skills (DLS) were measured with the Vineland Adaptive Behavior Scales (VABS), 1st (8) or 2nd edition (9), DLS Standard subscale was used to assess the participant's ability to perform personal and social tasks involved in everyday living (10). For the ASD group, the VABS consisted of a structured interview of a parent for child participants or an informant, when available, for adult participants.

#### Neuroimaging Data Acquisition and Preprocessing

Results included in this manuscript come from preprocessing performed using fMRIPrep (v23.1.3) (11), which is based on Nipype (v1.8.6) (12,13). Additionally, the ABIDE dataset offers preprocessed resting state fMRI data for participants scanned for ABIDE I that was manually inspected by three independent raters. These raters individually examined the quality of the raw anatomical and functional data, as well as the derivatives, and gave a pass/fail rating.

##### Anatomical Data Preprocessing

A total of 1 T1-weighted (T1w) images were found within the input BIDS dataset. The T1-weighted (T1w) image was corrected for intensity non-uniformity (INU) with `N4BiasFieldCorrection` (14), distributed with ANTs (version unknown) (15), and used as T1w-reference throughout the workflow. The T1w-reference was then skull-stripped with a Nipype implementation of the antsBrainExtraction.sh workflow (from ANTs), using OASIS30ANTs as target template. Brain tissue segmentation of cerebrospinal fluid (CSF), white-matter (WM) and gray-matter (GM) was performed on the brain-extracted T1w using `fast` (16). Brain surfaces were reconstructed using `recon-all` (FreeSurfer v7.3.2; RRID:SCR_001847) (17), and the brain mask estimated previously was refined with a custom variation of the method to reconcile ANTs-derived and FreeSurfer-derived segmentations of the cortical gray-matter of Mindboggle (Mindboggle; RRID:SCR_002438) (18). Volume-based spatial normalization to one standard space (MNI152NLin2009cAsym) was performed through nonlinear registration with antsRegistration (ANTs; version unknown), using brain-extracted versions of both T1w reference and the T1w template. The following template was selected for spatial normalization and accessed with TemplateFlow (v23.0.0) (19): ICBM 152 Nonlinear Asymmetrical template version 2009c (20).

##### Functional Data Preprocessing

For each of the 1 BOLD runs found per subject (across all tasks and sessions), the following preprocessing was performed. First, a reference volume and its skull-stripped version were generated using a custom methodology of fMRIPrep. Head-motion parameters with respect to the BOLD reference (transformation matrices, and six corresponding rotation and translation parameters) are estimated before any spatiotemporal filtering using FSL's `mcflirt` (21). BOLD runs were slice-time corrected to 0.714s (0.5 of slice acquisition range 0s-1.43s) using `3dTshift` from AFNI (RRID:SCR_005927) (22). The BOLD time-series (including slice-timing correction when applied) were resampled onto their original, native space by applying the transforms to correct for head-motion. These resampled BOLD time-series will be referred to as preprocessed BOLD in original space, or just preprocessed BOLD. The BOLD reference was then co-registered to the T1w reference using `bbregister` (FreeSurfer) which implements boundary-based registration (23). Co-registration was configured with six degrees of freedom. Several confounding time-series were calculated based on the preprocessed BOLD: framewise displacement (FD), DVARS and three region-wise global signals. FD was computed using two formulations following Power (absolute sum of relative motions, Power et al. (2014) and Jenkinson (relative root mean square displacement between affines, Jenkinson et al. (2002). FD and DVARS are calculated for each functional run, both using their implementations in Nipype (following the definitions by Power et al.). The three global signals are extracted within the CSF, the WM, and the whole-brain masks. Additionally, a set of physiological regressors were extracted to allow for component-based noise correction (25). Principal components are estimated after high-pass filtering the preprocessed BOLD time-series (using a discrete cosine filter with 128s cut-off) for the two CompCor variants: temporal (tCompCor) and anatomical (aCompCor). tCompCor components are then calculated from the top 2% variable voxels within the brain mask. For aCompCor, three probabilistic masks (CSF, WM and combined CSF+WM) are generated in anatomical space. The implementation differs from that of Behzadi et al. in that instead of eroding the masks by 2 pixels on BOLD space, a mask of pixels that likely contain a volume fraction of GM is subtracted from the aCompCor masks. This mask is obtained by dilating a GM mask extracted from the FreeSurfer's `aseg` segmentation, and it ensures components are not extracted from voxels containing a minimal fraction of GM. Finally, these masks are resampled into BOLD space and binarized by thresholding at 0.99 (as in the original implementation). Components are also calculated separately within the WM and CSF masks. For each CompCor decomposition, the k components with the largest singular values are retained, such that the retained components' time series are sufficient to explain 50 percent of variance across the nuisance mask (CSF, WM, combined, or temporal). The remaining components are dropped from consideration. The head-motion estimates calculated in the correction step were also placed within the corresponding confounds file. The confounded time series derived from head motion estimates and global signals were expanded with the inclusion of temporal derivatives and quadratic terms for each (26). Frames that exceeded a threshold of 0.5 mm FD or 1.5 standardized DVARS were annotated as motion outliers. Additional nuisance timeseries are calculated by means of principal components analysis of the signal found within a thin band (crown) of voxels around the edge of the brain, as proposed by (27). The BOLD time-series were resampled into standard space, generating a preprocessed BOLD run in MNI152NLin2009cAsym space. First, a reference volume and its skull-stripped version were generated using a custom methodology of fMRIPrep. All resamplings can be performed with a single interpolation step by composing all the pertinent transformations (i.e. head-motion transform matrices, susceptibility distortion correction when available, and co-registrations to anatomical and output spaces). Gridded (volumetric) resamplings were performed using antsApplyTransforms (ANTs), configured with Lanczos interpolation to minimize the smoothing effects of other kernels (28). Non-gridded (surface) resamplings were performed using mri_vol2surf in FreeSurfer.

Many internal operations of fMRIPrep use Nilearn (v0.10.1) (29), mostly within the functional processing workflow. For more details of the pipeline, see the section corresponding to workflows in fMRIPrep's documentation.

Due to the 2mm isotropic functional resolution, it is likely that the hand-drawn habenula ROIs captured signal from surrounding regions; therefore, denoising the BOLD fMRI signal was an essential step in the preprocessing pipeline to ensure participant-level effects were not solely due to spurious common variance from partial voluming or noise effects. To reduce physiological and motion-related signal contamination in our BOLD fMRI signal, CompCor was implemented, which defined nuisance regressors from the principal components of WM and CSF signals, rather than just averaging the WM and CSF signals as done by other data-driven denoising methods (30). We regressed out these first three principal components from each WM and CSF, along with 12 motion parameters (6 rigid-body motion parameters and their first temporal derivatives).

#### Whole-Brain Habenula Functional Connectivity Analysis

##### Model Selection

To control for site-specific effects in the multisite ABIDE data, ComBat is a popular choice as it allows the random intercepts of sites to have varying variance over voxels. However, a significant disadvantage of ComBat is that it distorts this variance when covariates or sample sizes are imbalanced across sites (31). Within our ABIDE participant sample, site-related interaction effects were significant ($p < 0.001$) for all covariates, including diagnostic group, age, and sex. **Figures S1, S2, and S3** delineate these effects and highlight the heterogeneity of covariates across the 34 ABIDE sites. After examining these effects, we determined that ComBat was not an appropriate approach to adjust for site variance in our analyses as we did not want to impose the assumption that covariate effects were uniform across sites. This is critical for autism-specific datasets, where site-related variability can reflect substantive differences in demographic composition, recruitment and referral patterns, diagnostic practices, and underlying population characteristics, rather than nuisance variability attributable solely to scanner effects.

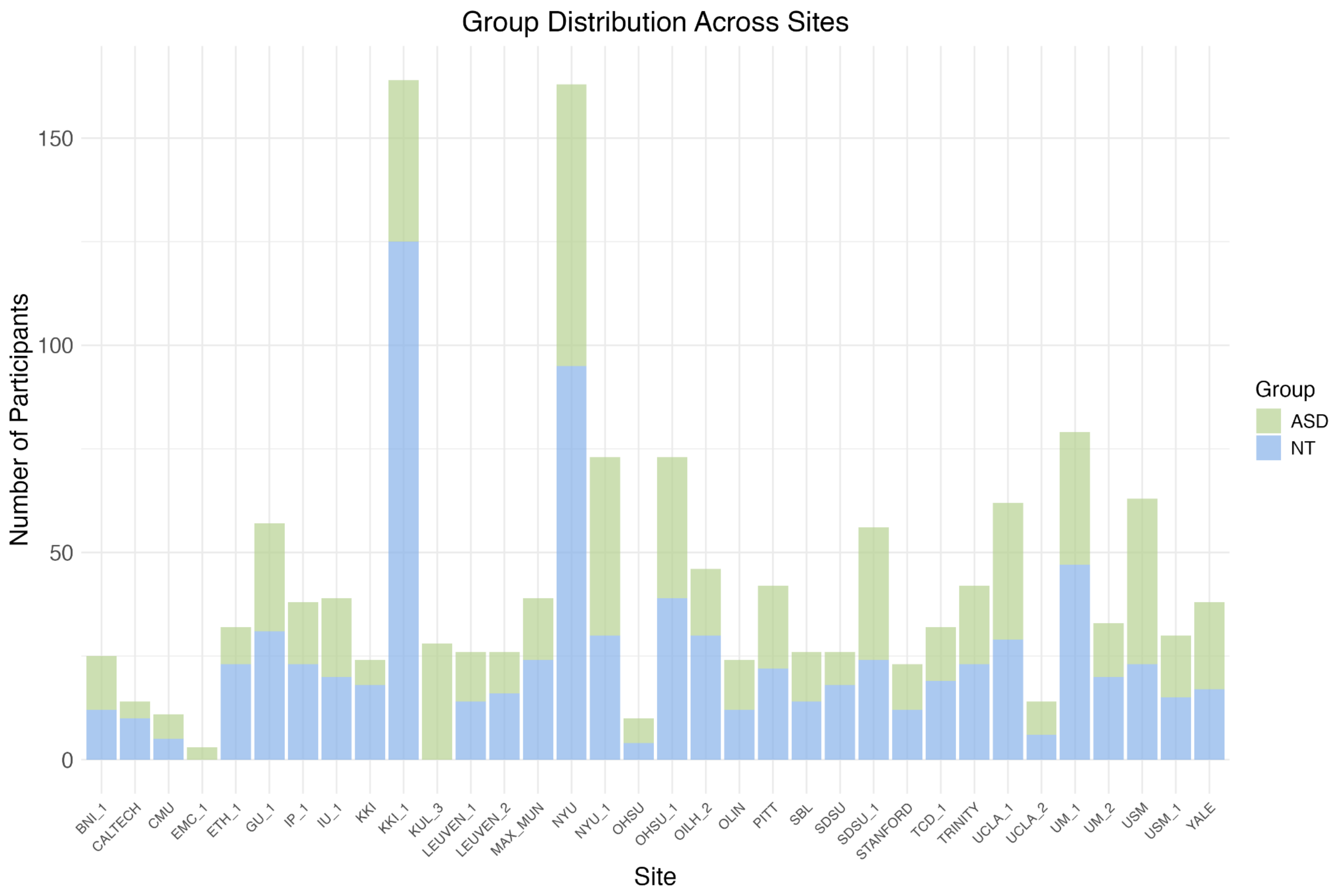


**Figure S1. Group Distribution Across Sites.** Participant composition by diagnostic group across the 34 ABIDE scanning sites, with stacked bars representing the number of individuals with autism spectrum disorder (ASD; green) and neurotypical (NT; blue) participants recruited at each location. The varying bar heights reflect differences in total sample size across sites (range: 3–164 participants), while the relative proportions illustrate marked heterogeneity in diagnostic composition, with ASD proportions ranging from 23.8% to 100%. A chi-square test confirmed significant non-independence between site and diagnostic group ($\chi^2 = 109.34$, $p = 4.14\times10^{-10}$), indicating that sites differed substantially in their relative recruitment of ASD versus NT participants.

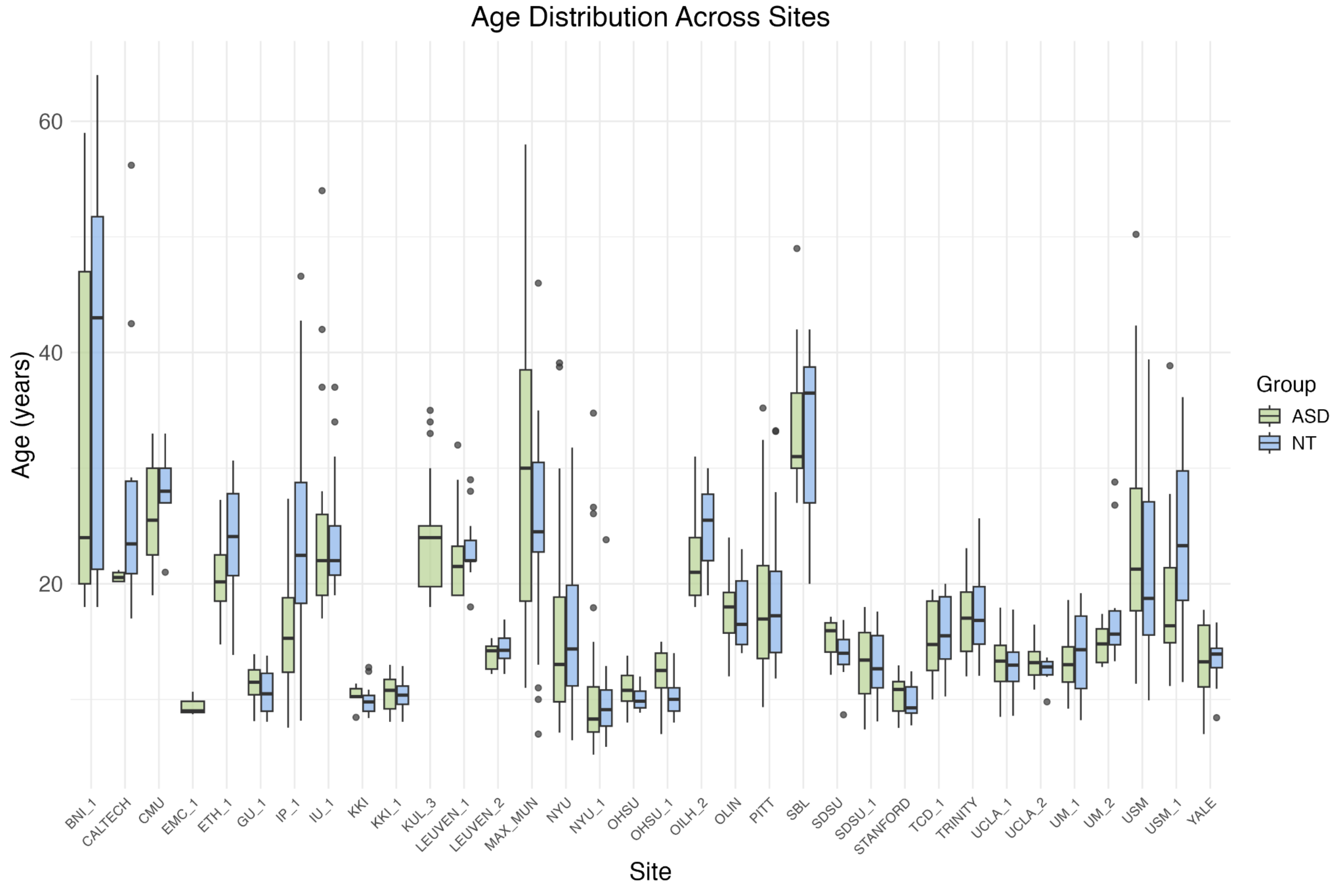


**Figure S2. Age Distribution Across Sites.** Distribution of participant ages across ABIDE scanning sites, shown separately for individuals with autism spectrum disorder (ASD; green) and neurotypical participants (NT; blue). A one-way ANOVA revealed significant differences in age distributions across sites ($F = 56.94$, $p < 2.13\times10^{-234}$), indicating substantial heterogeneity in the age composition of samples recruited from different scanning locations, with site-specific mean ages ranging from 9.5 to 36.5 years.

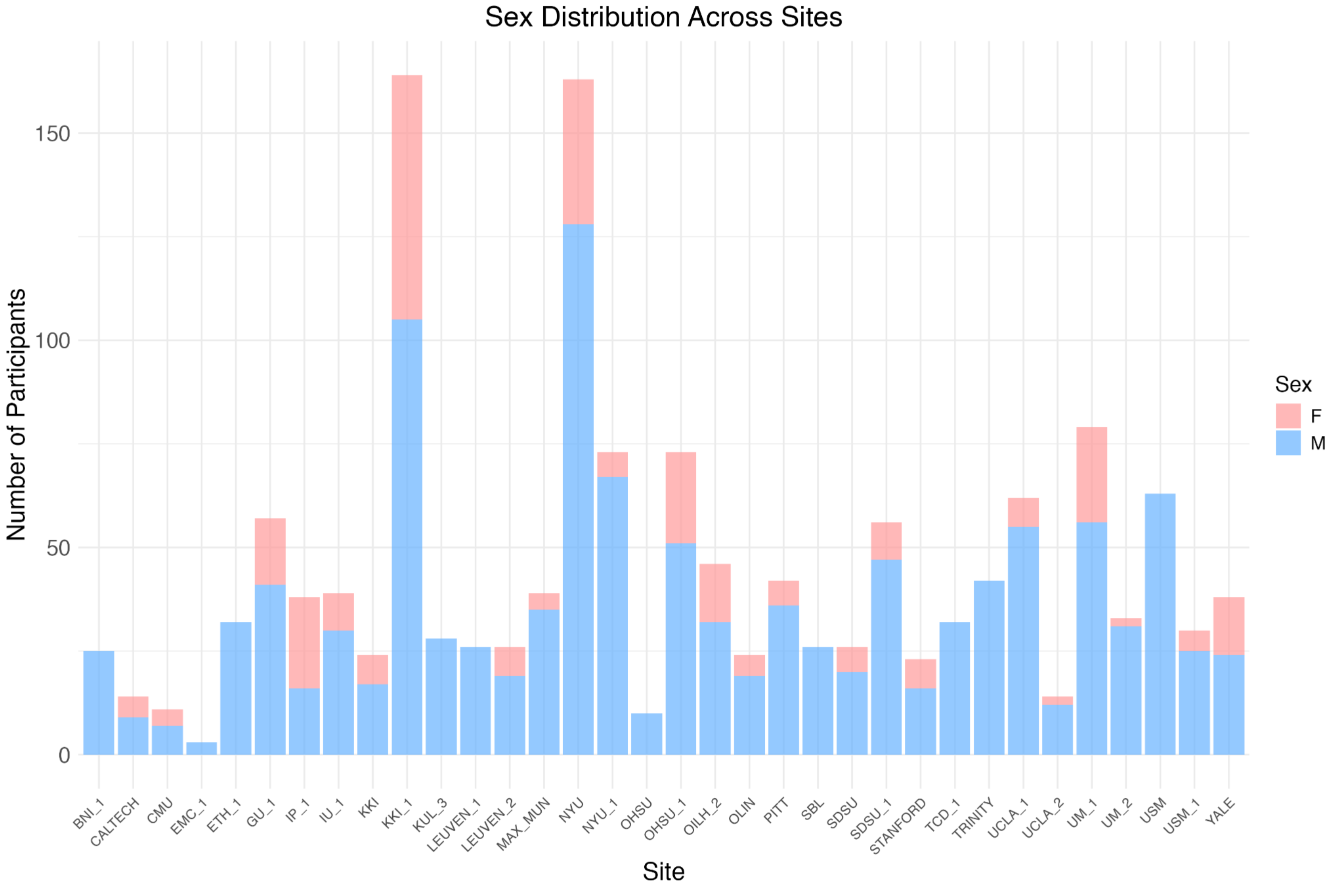


**Figure S3. Sex Distribution Across Sites.** Participant composition by biological sex across ABIDE scanning sites, with stacked bars representing the number of male (blue) and female (pink) participants recruited at each location. The varying bar heights reflect differences in total sample size across sites (range: 3–164 participants), while the relative proportions reveal considerable heterogeneity in sex composition, with male proportions ranging from 42.1% to 100% across sites. A chi-square test confirmed significant non-independence between site and sex ($\chi^2 = 178.79$, $p = 6.14\times10^{-22}$), demonstrating that scanning sites differed substantially in their sex composition.

Thus, given the site-related demographic heterogeneity in ABIDE, we prioritized a modeling framework that adjusted for site-related mean variability while explicitly estimating covariate and interaction effects of interest at the inferential stage. As such, we selected AFNI's mixed-effects modeling framework without harmonization. Following AFNI's `3dLMEr` documentation (https://afni.nimh.nih.gov/pub/dist/doc/program_help/3dLMEr.html), we accounted for scanning site variance by including it as a random effect in our linear-mixed effect model. Although AFNI's `3dLME` tool was built on the `nlme` R package, which allows for more complex approaches to model site differences, the tool itself does not support them. However, `3dLMEr` does have enhanced model specification for handling random-effects components, such as site.

AFNI's `3dLME` was also an important tool in appropriately mitigating the influence of partial voluming effects on our group-level findings. Specifically, we note that the implemented data-driven denoising method merely mitigated noise effects, rather than eliminating them entirely. Therefore, participant's whole-brain habenula connectivity maps may have reflected variability driven by various sources of noise, not true neural signals. For this reason, group-level analyses provided a critical framework for distinguishing reproducible population-level effects from idiosyncratic individual variability. The group-level rsFC analysis was performed by entering the whole-brain correlation maps from the participant-level rsFC analysis into a voxelwise linear mixed effects model using AFNI's `3dLMEr` (https://afni.nimh.nih.gov/pub/dist/doc/program_help/3dLMEr.html). This method identified voxels with shared temporal covariance (i.e., consistent patterns of correlated habenula BOLD fluctuations) across our 1,479 participants. Considering there are 1,082,025 GM voxels within the 2mm isotropic MNI-152 template, it is highly unlikely that non-systematic signal contamination due to partial voluming would show a strong and consistent effect across thousands of participants. Moreover, given our stringent statistical thresholds, with a voxelwise threshold of $p = 0.0001$ and cluster-level threshold of $p = 0.01$, any residual, non-systematic partial volume effects were even more unlikely to survive this correction (https://afni.nimh.nih.gov/pub/dist/doc/program_help/3dClustSim.html).

#### Supplemental Results

##### Whole-Brain Habenula Functional Connectivity Analysis

###### Group-Averaged Habenula Connectivity

Subcortically, positive habenula connectivity was most pronounced in a cluster of regions with peak connectivity in the thalamus. Habenular-thalamic connectivity spread throughout the medial dorsal nucleus, pulvinar, and the anterior nucleus. From the thalamus, habenula connectivity extended anteriorly to striatal regions encompassing the globus pallidus, putamen, and caudate, and inferiorly into midbrain regions such as the the substantia nigra (SN) and ventral tegmental area (VTA). Cortically, multiple clusters with strong positive habenula connectivity were identified in regions of the frontal and left parietal lobe. Within the prefrontal cortex, habenula connectivity peaked in the bilateral middle frontal gyri (BA 8/9) and extended to the superior frontal gyrus (BA 10). Within the left portion of the parietal cortex, the habenula had two clusters of peak connectivity in the postcentral gyrus (BA 7), and angular gyrus (BA 39) that expanded through the inferior and superior parietal lobes. Additionally, negative habenula connectivity was found with clusters of bilateral subcortical regions with stronger negative connectivity in the right cerebral regions. Negative habenula connectivity peaked in the anterior cingulate gyri (BA 32) and extended to the posterior cingulate gyri (BA 31).

###### Group Differences in Habenula Connectivity

Within the temporal lobe, habenula connectivity peaked in the bilateral middle temporal gyrus (MTG, BA 21) and subpeaked in the superior temporal gyrus (STG, BA 22).

#### Supplemental Discussion

##### Whole-Brain Habenula Functional Connectivity Analysis

###### Group-Averaged Habenula Connectivity

To provide additional evidence to assuage concerns regarding the habenula in standard-resolution datasets, we present a series of comparisons detailing the similarity of the connectivity maps we observed with those that have been previously published in the literature. In alignment with previously published habenula connectivity studies using high-resolution 3T (32) and 7T (33) fMRI, the strongest habenula connectivity effects were observed within the subcortical midbrain (**Fig. S4**) and (**Fig. S5**), respectively. Consistent across this prior work and our current study, the most prominent habenula connectivity was observed with the thalamus and extended through the caudate, putamen, and ventral tegmental area. Within the midbrain, we report strong positive habenula connectivity with the globus pallidus, nucleus accumbens, and substantia nigra, consistent with the results of the 3T study (32). In addition, across all studies, strong positive habenula connectivity was found with the posterior insula, hippocampus, as well as the anterior and dorsal ACC. In agreement with the 3T study (32), we report positive connectivity that extends through the anterior insula, as well as the sensory cortex, which includes the primary and secondary visual cortices and primary and early auditory regions (STG and STS). In terms of differences between our results and prior work, we note that our study examined a neurodiverse sample, while Ely et al. (2019) (32) and Torrisi et al. (2017) (33) examined neurotypical participants. Our observed subcortical connectivity patterns were more widespread, and less localized, compared to these prior studies. We identified positive connectivity within the septal nuclei and amygdala; notably, the 3T study reported negative amygdala connectivity. Additionally, while our study and the 7T study (33) identified positive connectivity with posterior ACC, the 3T study (32) did not. Interestingly, the 7T study (33) reported habenula connectivity with the left primary visual and auditory cortices, suggesting habenula involvement in left-lateralized sensory processing. However, not only do our group-averaged connectivity maps provide further evidence that habenula-sensory connectivity is bilateral, our group-difference results identified significantly increased connectivity with the bilateral MTG and STG among autistic individuals, which are involved in low-level sensory processing. Overall, our results are consistent with the habenula connectivity patterns previously reported in the literature and present novel findings that significantly expand current knowledge of the habenula's role in autism.

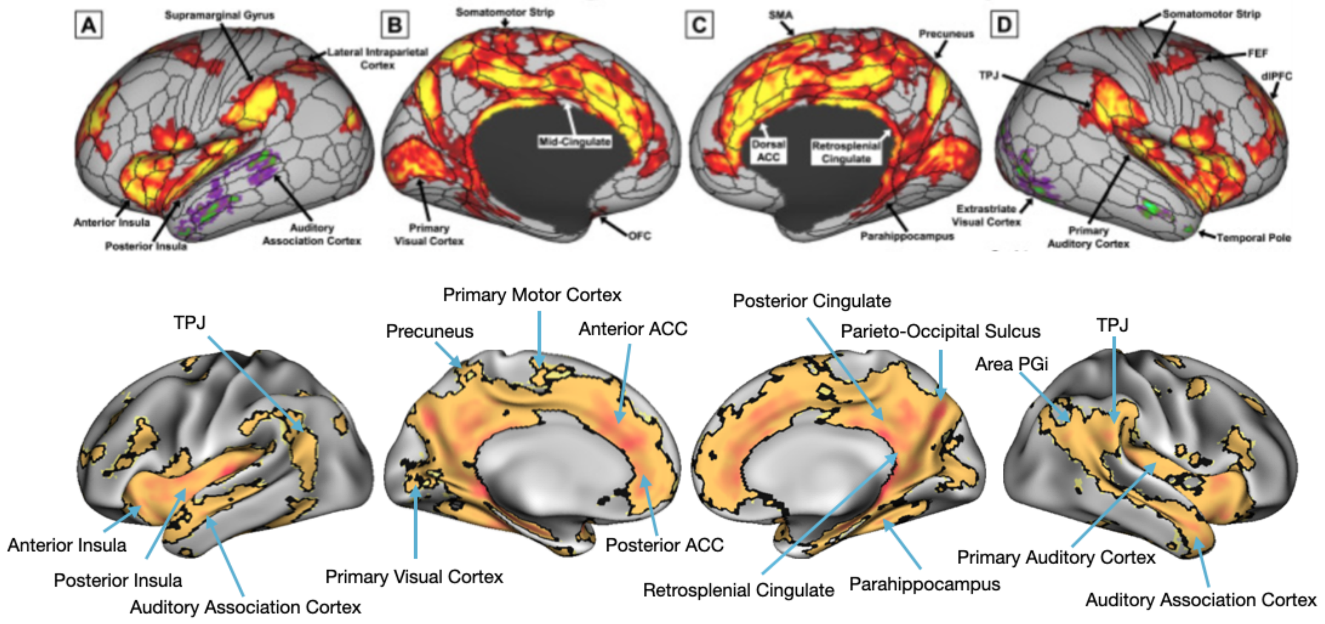


**Figure S4. Comparison of Habenula rsFC Results: Ely et al., 2019.** Whole-brain habenula rsFC surface maps published in a prior 3T fMRI study (Ely et al., 2019; top row) are consistent with the whole-brain habenula rsFC surface maps generated in the group-level analyses for the current study (bottom row).

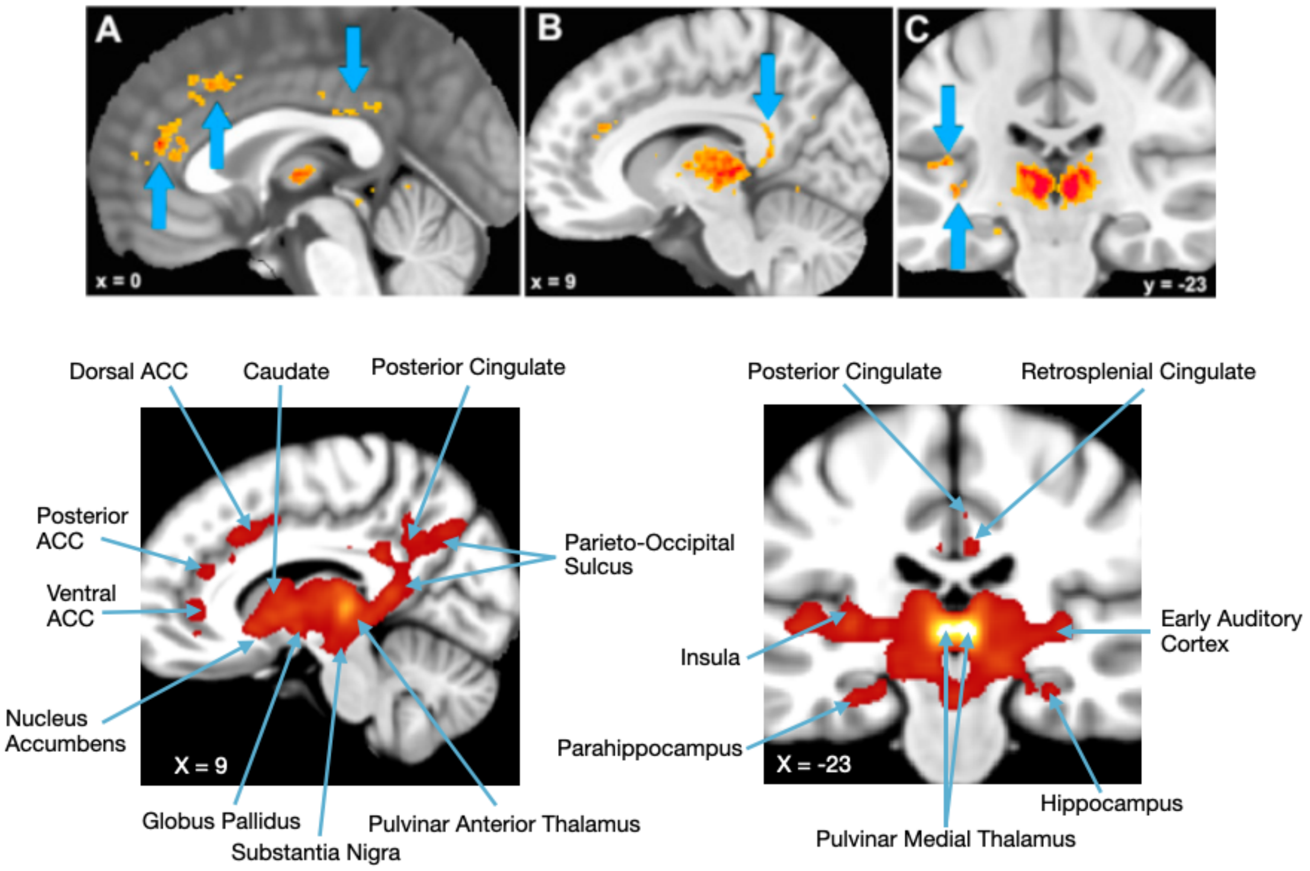


**Figure S5. Comparison of Habenula rsFC Results: Torrisi et al, 2017.** Whole-brain habenula rsFC volume maps published in a prior 7T fMRI study (Torrisi et al., 2017; top row) are consistent with the whole-brain habenula rsFC volume maps generated in the group-level analyses for the current study (bottom row).

14. Tustison NJ, Avants BB, Cook PA, Zheng Y, Egan A, Yushkevich PA, et al. N4ITK: improved N3

bias correction. IEEE Trans Med Imaging. 2010 Jun;29(6):1310–20. doi:10.1109/TMI.2010.2046908 PubMed PMID: 20378467; PubMed Central PMCID: PMC3071855.

27. Caballero-Gaudes C, Reynolds RC. Methods for cleaning the BOLD fMRI signal. NeuroImage. 2017

Jul 1;Cleaning up the fMRI time series: Mitigating noise with advanced acquisition and correction strategies154:128–49. doi:10.1016/j.neuroimage.2016.12.018
